# Anti-SARS-CoV-2 potential of *Cissampelos pareira* L. identified by Connectivity map-based analysis and in vitro studies

**DOI:** 10.1101/2021.06.11.448155

**Authors:** Madiha Haider, Vivek Anand, Dhwani Dholakia, M. Ghalib Enayathullah, Yash Parekh, Sushma Ram, Surekha Kumari, Anmol, Kiran Kumar Bokara, Upendra Sharma, Bhavana Prasher, Mitali Mukerji

**Author notes:** Corresponding Authors Genomics & molecular medicine, CSIR-Institute of Genomics and Integrative Biology, Delhi, India-110007, **Email:**, **Email:****. Department of Bioscience & Bioengineering, Indian Institute of Technology Jodhpur, NH 62, Karwar, Rajasthan 342037.

## Abstract

**Background:** Viral infections have a history of abrupt and severe eruptions through the years in the form of pandemics. And yet, definitive therapies or preventive measures are not present.

**Purpose:** Herbal medicines have been a source of various antiviral compounds. An accelerated repurposing potential of antiviral herbs can provide usable drugs and identify druggable targets. In this study, we dissect the anti-coronavirus activity of *Cissampelos pareira* L (*Cipa*). using an integrative approach.

**Methods:** We analysed the signature similarities between predicted antiviral agents and *Cipa* using the connectivity map (https://clue.io/). Next, we tested the anti-SARS-COV-2 activity of *Cipa in vitro*. A three-way comparative analysis of *Cipa* transcriptome, COVID-19 BALF transcriptome and CMAP signatures of small compounds was also performed.

**Results:** Several predicted antivirals showed a high positive connectivity score with *Cipa* such as apcidin, emetine, homoharringtonine etc. We also observed 98% inhibition of SARS-COV-2 replication in infected Vero cell cultures with the whole extract. Some of its prominent pure constituents e.g pareirarine, cissamine, magnoflorine exhibited 40-80% inhibition. Comparison of genes between BALF and *Cipa* showed an enrichment of biological processes like transcription regulation and response to lipids, to be downregulated in *Cipa* while being upregulated in COVID-19. CMAP also showed that Triciribine, torin-1 and VU-0365114-2 had positive connectivity with BALF 1 and 2, and negative connectivity with *Cipa*.

## Introduction

SARS-CoV-2, the severe acquired respiratory syndrome agent coronavirus 2, has taken many lives in the past year and is continuing to create an unsafe environment. Along with numerous mutations, fast transmission and a wide range of symptoms, lack of a definite therapeutic intervention has made this virus all the more deadly. Many studies have been conducted in order to recognize small compound therapeutics effective against SARS-CoV-2. Interestingly, some estrogen receptor modulators and protein synthesis inhibitors having potential antiviral against SARS-CoV2 have also been identified (1). It has been shown that *ESR1* as a drug target can modulate certain coronavirus associated genes. A group recently demonstrated the downregulation of *ACE2* by estrogen (2).

*Cissampelos pareira* L. is a commonly used hormone modulator which is used to treat reproductive disorders and fever. It has also been found to inhibit three serotypes of dengue (3) and its effect on various hormones has also been evidenced (4). In a previous study we have observed that *Cipa* can act as both, a protein synthesis inhibitor and an estrogen receptor inhibitor (5). Many of the drugs positively connected with *Cipa* have been reported to be a potential antiviral agent. Since there were several overlaps between the therapeutics predicted to be effective against SARS-CoV-2 and our formulation, we decided explore the repurposing potential of *Cipa* for this current pandemic.

## Material and Methods

### Transcriptome meta-analysis

We obtained the raw RNA sequencing data from SARS-CoV-2 patients Broncho-alveolar lavage fluid (BALF) 1 and 2 from the recent publications and analyzed them inhouse (supplementary information 1.1). The gene expression data for *Cissampelos pareira* L. was taken from (5), which can be accessed at GSE156445.

### Functional enrichment and connectivity map of the differentially expressed genes

The differentially expressed genes were analyzed for functional enrichment using enrichr (6) and for similar signatures using clue.io (7). The results were then compared with *Cipa* to find intersections between gene ontologies, enriched gene sets, and connectivity map perturbations between upregulated genes of BALF 1 and 2 and downregulated genes of *Cipa* and between downregulated genes of BALF 1 and 2 and upregulated genes of *Cipa*. Enrichments were also done for the genes whose knockdown signatures score >90 connectivity with *Cipa* and then compared with the upregulated processes and gene sets in SARS-CoV-2.

### Collection of plant material and preparation of extract for *in vitro* inhibition assays

Whole plant of *Cissampelos pareira* L was collected from the Palampur, HP, India (alt. 1350 m). The identification of the plant material was done by a taxonomy expert in CSIR-IHBT, Palampur and a voucher specimen (no. PLP16688) was deposited in the herbarium of CSIR-IHBT, Palampur, HP-176,061, India. The plant has been obtained and isolated as per CSIR, India guidelines (details for extract preparation in supplementary information 1.2). The pure molecules from roots of the plant were isolated as reported recently (8).

### Cell culture, viral infection and drug treatment for inhibition of SARS-COV-2 by *Cipa*

The effect of PE50 and PER was tested against the SARS-CoV2 (ASTM, 2015) in a 96-well tissue culture plates that was seeded with Vero Cells 24 h prior to infection with SARS-CoV2 (Indian/a3i clade/2020 isolate) in BSL3 facility. After treatment, RNA was isolated using MagMAXTM Viral/Pathogen Extraction Kit (Applied Biosystems, Thermofisher) according to the manufacturer’s instructions. The details of the experiment and RNA isolation protocol are given in the supplementary information 1.3.

### TaqMan Real-time RT-PCR assay for Detection of SARS-CoV-2

**T**he detection of genes specific to SARS-CoV2 was done using COVID-19 RT-qPCR Detection Kit (Fosun 2019-nCoV qPCR, Shanghai Fosun Long March Medical Science Co. Ltd.) according to the manufacturer’s instructions (details in supplementary information 1.4). The calculations for the relative viral RNA content and log reduced viral particles was calculated using the linear regression equation obtained using the RNA extracted from the known viral particles by RT-qPCR, using N, E and ORF1ab genes specific to SARS CoV2 virus. from the test sample (9).

## Results

### Cissampelos pareira L. shows high positive connectivity with small compounds predicted to inhibit SARS-CoV-2

To assess which among the numerous small compounds predicted to have antiviral potential against SARS-CoV2 in various studies might have similar signatures among the same cell lines as *Cipa*, we queried the connectivity map. We observed 7 small compounds to possess high signature similarity with *Cipa*. These include emetine (99.61), anisomycin (99.58), cycloheximide (99.86), homoharringtonine (99.51), apcidin (86.32), ruxolitinib (91.54), and sirolimus (95.45). These small compounds were predicted using different methods in 4 different studies (Table 1).

**Table 1.**
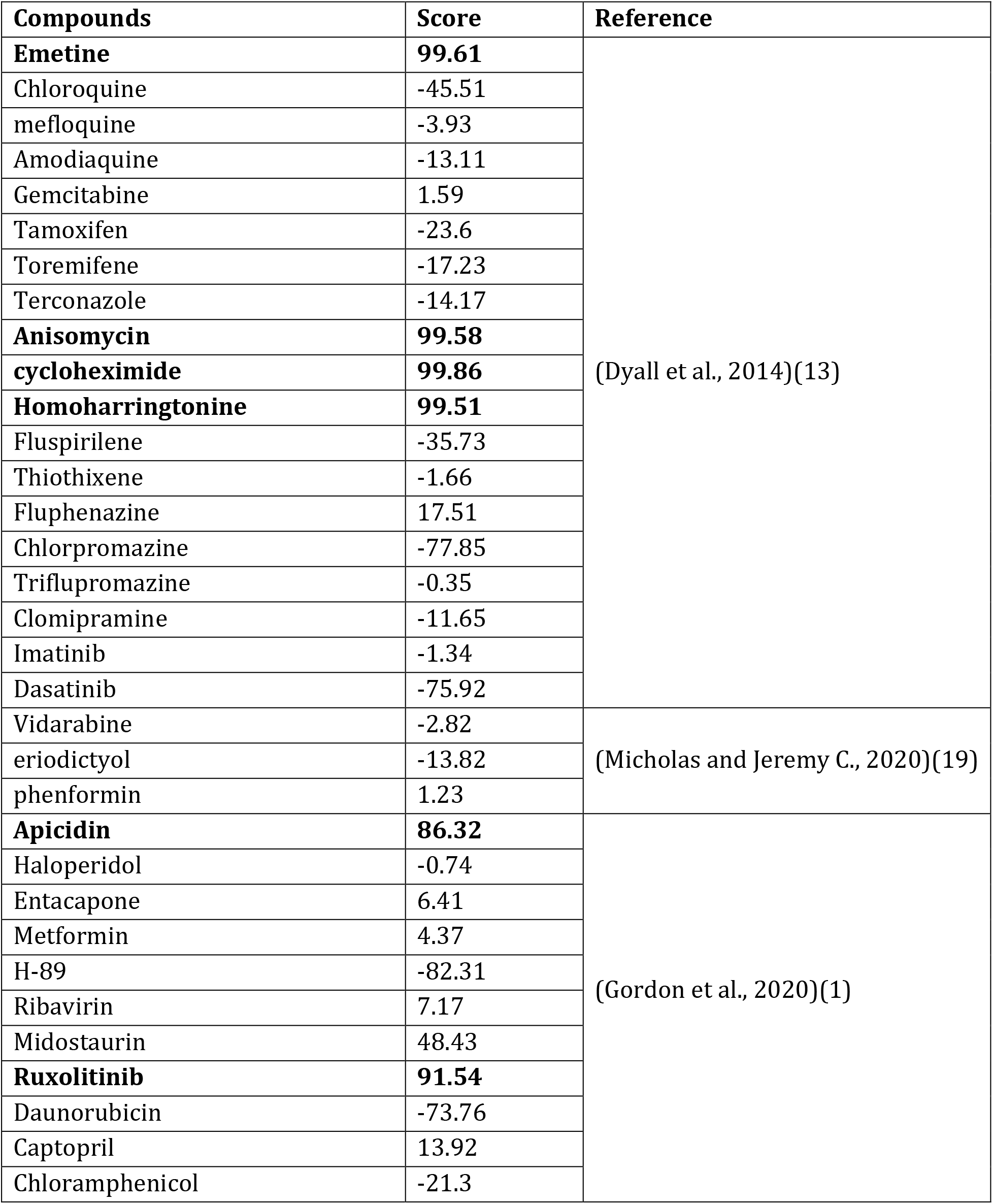
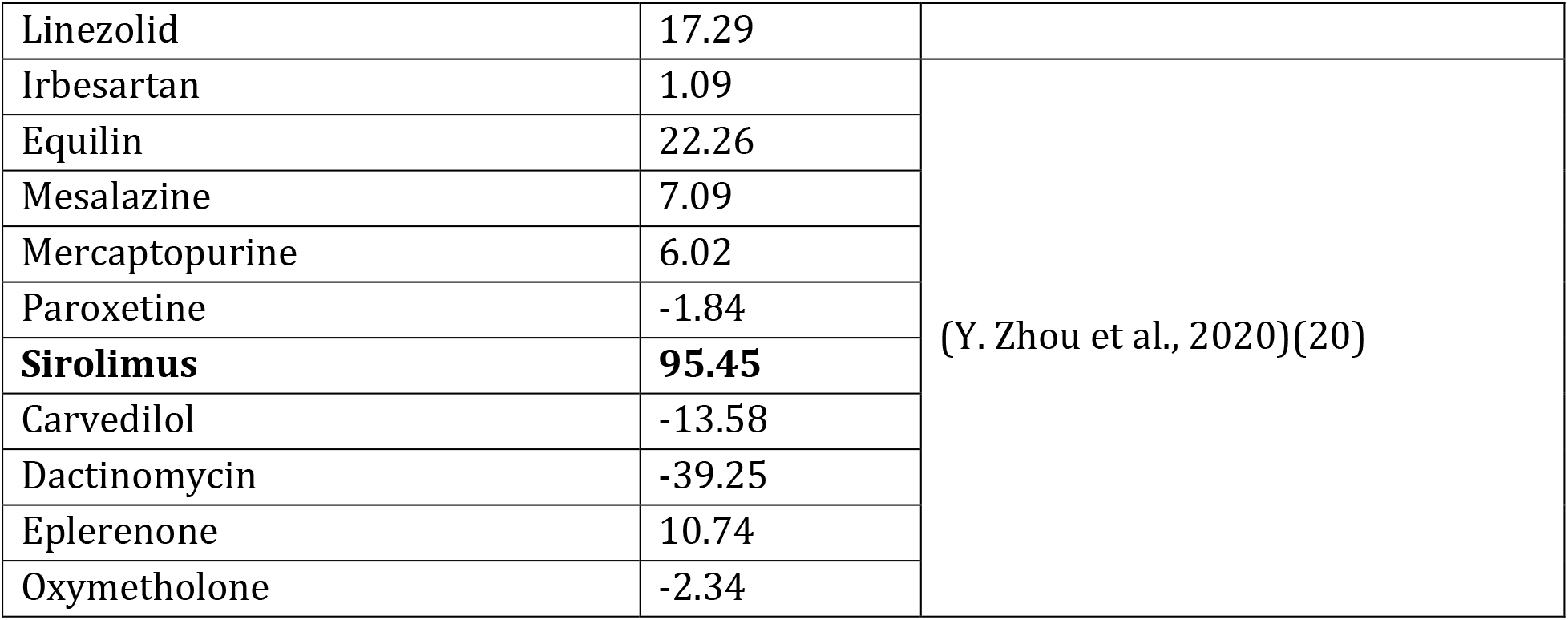
Connectivity scores of small compounds with repurposing potential against SARS-CoV-2 as predicted by various studies.

### Cipa whole extract and single molecule constituents can inhibit SARS-COV-2 in vitro

Since metanalysis highlighted an inhibitory potential of *Cipa* against SARS-COV-2, we tested the effect of whole plant and root extracts of *Cipa* in Vero cell culture assays infected with SARS-CoV-2. The relative viral RNA (%) was calculated by considering the values averaged from N (Nucleoprotein), ORF1ab (19 non-structural proteins, NSP1-16), and E (Envelope) viral genes. The whole plant aqueous extract showed a definite antiviral activity, evidenced by decreased relative viral RNA content with a reduction by 57% at 100μg/ml where the viral particle number reduced from 10^5.9^ to 10^5.6^ (Figure 1A-C).

**Figure 1:**
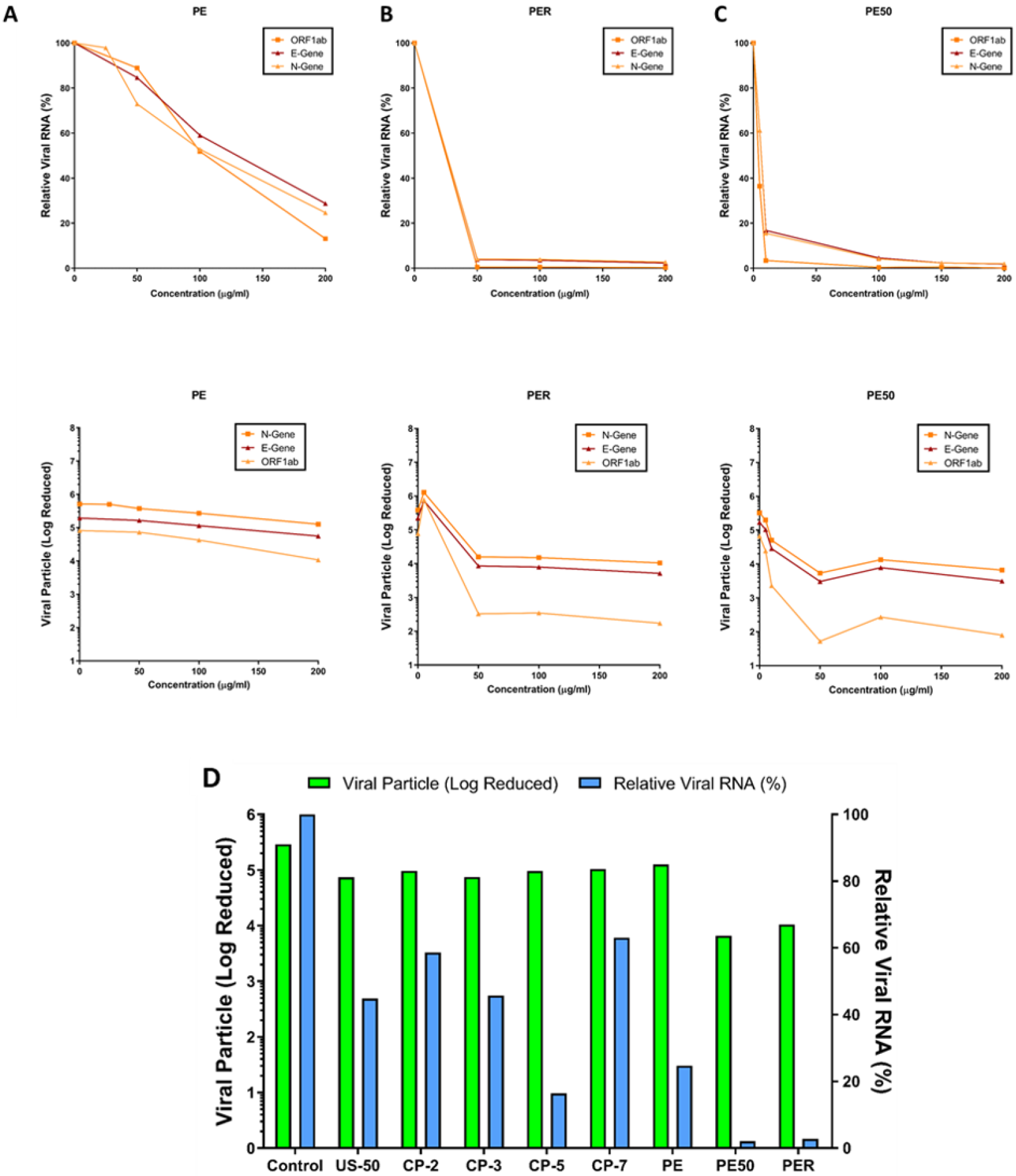
Inhibition of SARS-CoV-2 in vitro by Cipa whole extract and its constituents: Relative viral RNA % and Log reduction in viral particles in vero cells upon treatment at 50, 100, 150 and 200μg of A) whole plant aqueous extract (PE), B) root extract (PER) and C) hydro-alcoholic extracts (PE50) of Cipa. D) Sars-cov-2 viral titers inhibition by Cipa constituents CP-2 Salutaridine, CP-3 Cissamine, CP-5 pareirarine, CP-7 Magnoflorine, PE aqueous extract, PE50 50% hydroalcoholic extract and PER root extract.

Next, we wanted to see whether the *Cipa* pure constituents can inhibit SARS-COV-2. The total alkaloid content in the extract was 46.4 mg/g with cissamine (18.6 mg/g) being the major one, followed by magnoflorine (12.9 mg/g). The pure molecules namely hayatinin (US-50), salutaridine (US-DR-CP-2), cissamine (US-CP-3), pareirarine (US-CP-5), magnoflorine (US-CP-7), aqueous whole plant extract (PE), 50% hydroalcoholic whole plant extract (PE50) and 50% hydroalcoholic root extract (PER) were tested against SARS-CoV2 at 200 μM concentration showed relative viral RNA (%) to 44, 58, 45, 16, 63, 24, 2 and 2 respectively in comparison with the virus control (Figure 1D).

### Cipa transcriptome oppositely regulates several biological pathways compared to COVID-19 infected patient BALF transcriptome

We looked for the differentially expressed genes which were common between *Cipa* and BALF samples from two studies (10, 11), hereby referred to as BALF-1 and BALF-2 respectively. We observed that 39 genes were common between *Cipa* and BALF-1 and BALF-2, out of which 29 showed an opposite expression (Table S1). Individually, 134 genes were common between *Cipa* and BALF-1, and 174 genes common between *Cipa* and BALF-2. Upon functional enrichment analysis we observed that the genes enriched for regulation of vascular endothelial growth were upregulated in both BALF-1 and 2 datasets while being downregulated by *Cipa*. Similarly, while regulation of gene expression was upregulated by *Cipa* it was downregulated in BALF-1 and BALF-2 data sets (Figure 2A-C).

**Figure 2:**
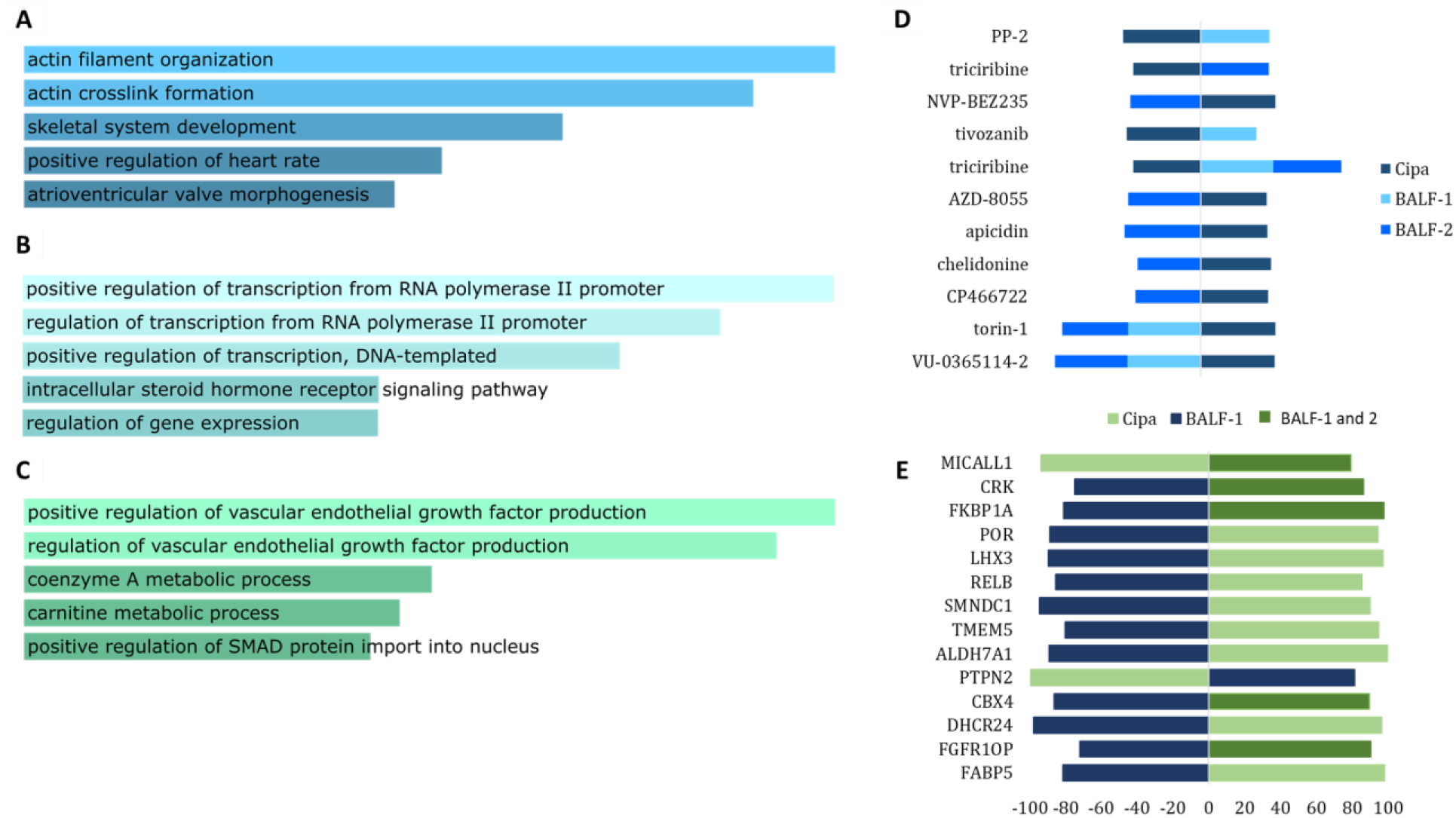
Comparative analysis of Cipa with BALF-1 and BALF-2 transcriptome: Functional enrichment of genes common between A) Cipa and BALF-1, B) Cipa and BALF-2 and C) Cipa, BALF-1 and BALF-2. D) Small compound signatures common between all three, the space on the left of the vertical axis indicates negative signatures while the one on the right indicates positive signatures. E) Genetic signatures common between all three, the space on the left of the vertical axis indicates negative signatures while the one on the right indicates positive signatures.

The connectivity map analysis of the gene signatures of BALF-1 and BALF-2 comparison with *Cipa* results revealed a number of small compounds that had high positive scores with *Cipa* and negative scores with either BALF transcriptomes. We observed several small compounds having opposite signature similarities between *Cipa* and BALF. Triciribine, torin-1 and VU-0365114-2 were three compounds with negative scores for *Cipa* and positive scores for both BALF 1 and 2 (Figure 2D). Also, knockdown signatures of a number of genes were found to exhibit opposing scores between Cipa and BALF transcriptomes. Some of these include, MICALL1, CRK, FKBP1A, CBX4, and FGR10P (Figure 2E).

## Discussion

SARS-COV-2 has shown a very diverse set of clinical presentations in various populations and among genders within the same population. It has been shown that estrogen can regulate the expression of ACE-2 receptors (2). Since *Cipa* appears to have *ESR1* modulatory effects (5), and has been shown to have antiviral potential (3), we hypothesized it may have inhibitory effect on the novel coronavirus. CMAP analysis of *Cipa* transcriptome signatures highlights several small compounds having been predicted to have inhibitory activity against SARS-CoV-2. Among these, emetine, homoharringtonine, and cycloheximide are known translation inhibitors. These have also been shown to inhibit Zika and Ebola (12), SARS and MERS (13), and Newcastle disease virus (14). Apcidin, an HDAC inhibitor has been predicted to inhibit SARS-CoV-2 in a recent study (15).

*In vitro* experiments for viral inhibition of SARS-COV-2, reveal that all whole plant extracts of *Cipa* (aqueous and alcoholic) could inhibit the virus at least up to 60%. Hydroalcoholic whole plant extract showed an inhibition of 98%. The single molecule constituents of Cipa could also inhibit the viral particles, with pareirarine showing the highest inhibition of 80%. This showed that *Cipa* does have the potential to inhibit SARS-CoV-2 virus *in vitro*. Interestingly, the highest inhibition is shown by the whole plant hydroalcoholic extract which comprises of various small compound constituents. This suggests a synergistic effect of the constituents towards viral inhibition.

We also found that the signatures of the transcriptomic changes in *Cipa* treated MCF7 cells and BALF from patients’ lungs, have interesting overlaps. Among the connected small compounds triciribine, torin-1 and VU-0365114-2, triciribine has been shown to inhibit Human Immunodeficiency virus sera types 1 and 2 (16). Another study has shown that VU-0365114-2, which is a muscarinic acetylcholine receptor M5 inhibitor, has repurposing potential against SARS-CoV-2 (17). While mTOR inhibitor torin-1 may modulate immune activity and enhance antiviral response, even against SARS-COV-2 (18).

## Conclusion

In summary, we report here a framework applicable for repurposing of herbal formulations using an integrated multi-pronged approach using transcriptome-based connectivity mapping, *in vitro* validation and conjoint analysis with disease signatures. We demonstrate the potential repurposing of *Cissampelos pareira* L for sars-cov-2 using this approach.

## Acknowledgements

The authors would like to thank Dr. Rakesh Mishra (director-CCMB) for facilitating and support for SARS-CoV-2 infection model. The authors acknowledge research fellowship support to MH (University Grants Commission), DD (Department of Biotechnology) SK (CSIR) and Anmol (DST-INSPIRE).

## Abbreviations

CMAP: Connectivity Map
BALF: Bronchoalveolar Lavage fluid
ESR1: Estrogen Receptor 1
ACE2: Angiotensin-converting enzyme 2
SARS: Severe Acute Respiratory Syndrome
MERS: Middle East respiratory syndrome
HDAC: Histone deacetylase

## References

1. D. E. Gordon, et al., A SARS-CoV-2 protein interaction map reveals targets for drug repurposing. Nature (2020) https://doi.org/10.1038/s41586-020-2286-9.

2. K. E. Stelzig, et al., Estrogen regulates the expression of SARS-CoV-2 receptor ACE2 in differentiated airway epithelial cells. Am. J. Physiol. Cell. Mol. Physiol. 318, L1280–L1281 (2020).

3. R. Sood, et al., Cissampelos pareira Linn: Natural Source of Potent Antiviral Activity against All Four Dengue Virus Serotypes. PLoS Negl. Trop. Dis. 9, 1–20 (2015).

4. M. Ganguly, M. Kr Borthakur, N. Devi, R. Mahanta, Antifertility activity of the methanolic leaf extract of Cissampelos pareira in female albino mice. J. Ethnopharmacol. 111, 688–691 (2007).

5. M. Haider, et al., Transcriptome analysis and connectivity mapping of *Cissampelos pareira* L. provides molecular links of ESR1 modulation to viral inhibition. bioRxiv, 2021.02.17.431579 (2021).

6. M. V Kuleshov, et al., Enrichr: a comprehensive gene set enrichment analysis web server 2016 update. Nucleic Acids Res. 44, W90–7 (2016).

7. A. Subramanian, et al., A Next Generation Connectivity Map: L1000 Platform and the First 1,000,000 Profiles. Cell (2017) https://doi.org/10.1016/j.cell.2017.10.049.

8. V. Bhatt, et al., Chemical profiling and quantification of potential active constituents responsible for the antiplasmodial activity of Cissampelos pareira. J. Ethnopharmacol. 262, 113185 (2020).

9. L. Caly, J. D. Druce, M. G. Catton, D. A. Jans, K. M. Wagstaff, The FDA-approved drug ivermectin inhibits the replication of SARS-CoV-2 in vitro. Antiviral Res. (2020) https://doi.org/10.1016/j.antiviral.2020.104787.

10. Y. Xiong, et al., Transcriptomic characteristics of bronchoalveolar lavage fluid and peripheral blood mononuclear cells in COVID-19 patients. Emerg. Microbes Infect. 9, 761–770 (2020).

11. Z. Zhou, et al., Heightened Innate Immune Responses in the Respiratory Tract of COVID-19 Patients. Cell Host Microbe 27, 883–890.e2 (2020).

12. S. Yang, et al., Emetine inhibits Zika and Ebola virus infections through two molecular mechanisms: inhibiting viral replication and decreasing viral entry. Cell Discov. 4, 31 (2018).

13. J. Dyall, et al., Repurposing of clinically developed drugs for treatment of Middle East respiratory syndrome coronavirus infection. Antimicrob. Agents Chemother. 58, 4885–4893 (2014).

14. H.-J. Dong, et al., The Natural Compound Homoharringtonine Presents Broad Antiviral Activity In Vitro and In Vivo. Viruses 10 (2018).

15. K. Liu, et al., Clinical HDAC Inhibitors Are Effective Drugs to Prevent the Entry of SARS-CoV2. ACS Pharmacol. Transl. Sci. 3, 1361–1370 (2020).

16. R. G. Ptak, et al., Inhibition of Human Immunodeficiency Virus Type 1 by Triciribine Involves the Accessory Protein Nef. Antimicrob. Agents Chemother. 54, 1512 LP–1519 (2010).

17. Z. Wang, et al., Identification of Repurposable Drugs and Adverse Drug Reactions for Various Courses of COVID-19 Based on Single-Cell RNA Sequencing Data. ArXiv, arXiv:2005.07856v2 (2020).

18. Y. Zheng, R. Li, S. Liu, Immunoregulation with mTOR inhibitors to prevent COVID-19 severity: A novel intervention strategy beyond vaccines and specific antiviral medicines. J. Med. Virol. 92, 1495–1500 (2020).

19. S. Micholas, S. Jeremy C., Repurposing Therapeutics for COVID-19: Supercomputer-Based Docking to the SARS-CoV-2 Viral Spike Protein and Viral Spike Protein-Human ACE2 Interface. chemRxiv (2020) https://doi.org/10.26434/chemrxiv.11871402.v4.

20. Y. Zhou, et al., Network-based drug repurposing for novel coronavirus 2019- nCoV/SARS-CoV-2. Cell Discov. 6, 14 (2020).

