## Supplementary Information for "Anti-SARS-CoV-2 potential of *Cissampelos pareira* L. identified by Connectivity map-based analysis and in vitro studies"

**Supplemetary Information for:**

[a]Genomics & molecular medicine, CSIR-Institute of Genomics and Integrative Biology, Delhi, India-110007, [b]Academy of Scientific and Innovative Research, Ghaziabad, Uttar Pradesh, India 201002, [c]CSIR-Center for Cellular and Molecular Biology, Hyderabad, Telangana, 500007, India, [d]Chemical Technology Division, CSIR-Institute of Himalayan Bioresource Technology, Palampur, Himachal Pradesh 176 061, [e]Centre of excellence for Applied developments of Ayurveda prakriti and genomics, CSIR's Ayurgenomics Unit TRISUTRA, CSIR-IGIB, India

*Corresponding Authors

Genomics & molecular medicine, CSIR-Institute of Genomics and Integrative Biology, Delhi, India-110007

** Current address: Department of Bioscience & Bioengineering, Indian Institute of Technology Jodhpur, NH 62, Karwar, Rajasthan 342037

**1. Supplementary methods:**

### 1.1 SARS-CoV-2 transcriptome meta-analysis:

RNA-Seq sample pre-processing:
Created annotated transcripts from human genome (GRCh37) and GTF file (gencode v19) using gffread (v0.12.1) and further indexing the transcripts using Salmon index (v1.3.0).
salmon index --threads $cpus -t $fasta --gencode -i salmon_index 2

Raw reads were mapped and quantified using salmon quant with options --validateMappings (**improve quantification accuracy**), --seqBias (**correct sequence-specific bias**), --useVBOpt (**use Bayesian EM algorithm**), --gcBias (**correct GC bias**).
salmon quant --validateMappings --seqBias --useVBOpt --gcBias --geneMap $gtf --threads $cpus --libType=A --index $salmon_index -1 $reads1.fq -2 $reads2.fq --writeUnmappedNames -o $sample

Salmon gives transcript level abundance for each sample and later converted into gene-level expression using tximport (v1.16.1). Before moving to differential gene expression, Surrogate Variable Analysis (v3.36.0) in an unsupervised setting to correct the batch effect in the samples considering the sequencing machine bias in both groups.

### 1.2 Preparation of *Cissampelos pareira* L. extracts and isolation of small compounds:

**Chemicals and reagents:** For UPLC analysis formic acid was obtained from S. D. Fine Chemicals Ltd. (Mumbai, India) whereas methanol and water (LC grade) were purchased from J. T. Baker (Mallinckrodt Baker Inc., St. Louis, MO, USA).
**Preparation of Extracts and Isolation of Pure Compounds:** The collected plant material was dried using shade drying method. The air dried whole plant (root, stem and leaves) material (230 g) was extracted thrice with ethanol: water (50: 50) by percolation. The percolate was evaporated to dryness in rotary evaporator at temperature 50 °C to obtain 25.0 g (10.8% w/w) crude extract (USCPWP-PE-50). The air dried roots (950 g) were extracted thrice with ethanol: water (80: 20) by percolation at room temperature. The percolate was evaporated in rotary evaporator at temperature 50 °C to yield 98.0 g (10.3 % w/w) crude extract (USCPR-PE-R) (Fig. S1). The whole plant aqueous extract was obtained commercially from a GMP certified manufacturer.
**Quantitative analysis of the samples:** The marker compounds in crude extract (USCPWP-PE-50) were quantified by the UPLC-DAD method reported recently (Bhatt et al., 2020).

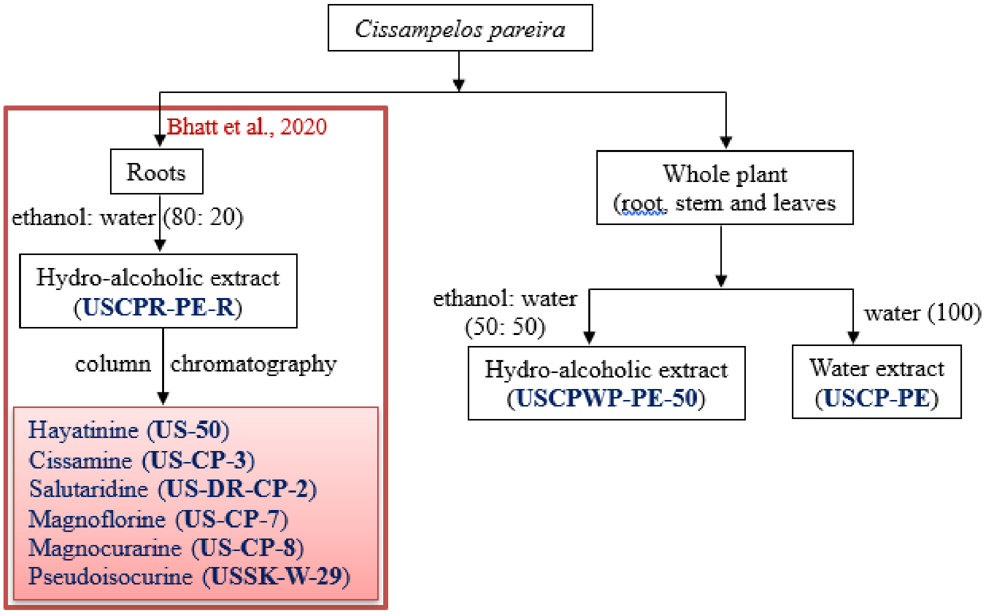


**Fig. S1: Flow chart of preparation of extracts, and isolation of pure molecules from Cissampelos pareira.**

#### 1.3 Cell culture, viral infection and drug treatment for inhibition of SARS-COV-2 by Cipa:

The cells were maintained in Dulbecco Minimum Essential Medium (DMEM) (Gibco) containing 10% Fetal Bovine Serum (FBS) (Gibco) at 37°C, 5% CO2. Initially the compounds were dissolved in organic and aqueous solvents based on the requirements, and stocks were made. The concentrations (200, 100, 50, 10, 5 (µg/mL), were made using the DMEM media. Briefly, the cells were pre-incubated with the compounds (PE-50 & R) for 2 hours, for each concentration in triplicate. Later, the virus inoculum (at a 0.1 MOI) was added to the cells for 3 hours in presence of the respective dilutions of compounds made in Basal medium only. Post-infection, viral inoculum was replaced with fresh media containing 10% FBS and the experiment is continued in the presence of different dilutions of compounds for 72 hours. After 72 hours, cell supernatant was collected and spun for 10 min at 6,000 g to remove debris and the supernatant was transferred to fresh collection tubes for further analysis.

Isolation of Viral RNA:
RNA was isolated from 200 µL of the supernatants using the The viral supernatants from the test groups were added into the deep well plate (KingFisherTM Thermo Scientific) along with a lysis buffer containing the following components - MagMAXTM Viral/Pathogen Binding Solution (260µL); MVP-II Binding Beads (10 µL); MagMAXTM Viral /Pathogen Proteinase-K of (5 µL) respectively. RNA extraction was performed using KingFisher Flex (version 1.01, Thermo Scientific) by following manufacturer's instructions. The eluted RNA was immediately stored in -80$^{0}$C until further use.

### 1.4 TaqMan Real-time RT-PCR assay for Detection of SARS-CoV-2:

The kit detects Envelope gene (E; ROX labelled), Nucleocapsid gene (N- JOE labelled) and open reading frame 1ab (ORF1ab, FAM labelled) specific to SARS-CoV2 for detection and amplification of the cDNA. Briefly, RT-qPCR assays were performed on a Quant studio Q5 (Thermo fisher). A 10 µL of Negative Control, 10 µL of Positive Control (positive and negative controls were provided by kit), and 10 µL of extracted RNA from samples were added in different PCR reaction tubes. The contents were centrifuged at low speed. The cycling conditions are: Step 1: 50°C for 15 minutes, 1 cycle; Step 2: 95°C for 3 minutes, 1 cycle; Step 3: 95°C for 5 seconds to 60°C for 40 seconds, 5 cycles; Step 4: 95°C for 5 seconds to 60°C for 40 seconds, 40 cycles. The signals of FAM, JOE, ROX and CY5 (internal reference) fluorescence channels were collected at 60°C. SARS-CoV-2 cDNA (Ct~28) was used as a positive control. The log viral particles and a semi–log graph was plotted using Graph Pad Prism 5 software (ver 5.03).

### 2. Supplementary table:

**Table S1: 39 genes common between Cipa, BALF-1 and BALF-2**

| | Cipa | BALF-1 | BALF-2 |
|---|---|---|---|
| *AGR3* | -1.38746 | -1.18033 | -4.78508 |
| *MSMB* | -1.33061 | 1.070106 | -5.04154 |
| *ACLY* | -1.30041 | 0.295665 | 3.559965 |
| *PBX1* | -1.24767 | 1.487727 | 3.468367 |
| *PDIA4* | -1.24474 | 0.117456 | 6.354455 |
| *CROT* | -1.20234 | -0.83981 | 3.680022 |
| *C1orf21* | -1.17113 | -1.92896 | 2.874781 |
| *SCAI* | -1.1355 | -0.72477 | 4.309578 |
| *BCL9* | -1.1204 | 0.064614 | 4.146401 |
| *PORCN* | -1.08959 | -0.59788 | 3.219026 |
| *EVL* | -1.08953 | -1.47847 | 7.791948 |
| *TTLL5* | -1.0657 | -0.10307 | 2.754773 |

| | | | |
|---|---|---|---|
| *ETNK2* | -1.04426 | -2.07727 | 5.625841 |
| *TGFB1* | -1.04116 | -0.93531 | 2.21809 |
| *ANTXR1* | -1.03224 | -1.01169 | 3.570147 |
| *TET1* | -1.032 | -0.90988 | 5.321577 |
| *NUBPL* | -1.01577 | -0.12047 | 4.078702 |
| *SBK1* | -1.01475 | -2.39136 | 2.946711 |
| *RBPMS* | -1.00924 | -0.88967 | 2.967272 |
| *KIAA1217* | 1.012205 | -0.73144 | -2.07887 |
| *CBR3-AS1* | 1.030189 | -0.18835 | 3.847575 |
| *TBC1D12* | 1.037079 | 1.18878 | 4.152344 |
| *RANBP6* | 1.059566 | -0.34571 | 4.111845 |
| *STOM* | 1.080323 | 0.232034 | 3.092429 |
| *SLC22A5* | 1.090643 | -0.38776 | 5.721312 |
| *CNKSR3* | 1.116294 | 0.0481 | 4.422747 |
| *PLD1* | 1.122033 | 1.093094 | 2.99798 |
| *ZNF461* | 1.189381 | -0.68859 | 4.836408 |
| *LMO7* | 1.22905 | -1.23364 | -2.8111 |
| *CHD2* | 1.277242 | -0.59575 | 2.136096 |
| *EIF2AK3* | 1.341297 | 0.10631 | 3.506099 |
| *MYO9A* | 1.616131 | -0.78824 | 4.192263 |
| *PSPC1* | 1.624655 | -0.5562 | 4.292772 |
| *NEDD4* | 1.711919 | -0.95851 | 2.477732 |
| *ZNF462* | 1.867743 | -3.00318 | 6.111656 |
| *MYO1B* | 2.048216 | 0.553853 | -2.9671 |
| *TAGLN* | 2.238096 | -0.80487 | 6.711373 |
| *CYP1B1* | 2.547549 | 1.94272 | -4.75062 |
| *RND3* | 3.153523 | -1.67392 | -3.60436 |
